# Environmental versus litter traits as drivers of microbial decomposer functions

**DOI:** 10.1101/2024.07.26.605152

**Authors:** Svenja C. Stock, Rafaella Canessa, Liesbeth van den Brink, Lohengrin A. Cavieres, Andreas Kappler, Carolina Merino Guzmán, Alfredo Saldaña, Thomas Scholten, Andreas H. Schweiger, Kira Rehfeld, Harald Neidhardt, Yvonne Oelmann, Katja Tielbörger, Maaike Y. Bader, Todd A. Ehlers, Michaela A. Dippold

## Abstract

Plant litter decomposition is a key ecosystem process with significant implications for global carbon cycling, soil fertility and plant productivity. Given that microbial decomposers are the main players in the decomposition process, it is surprising how little is known about their functional diversity in different habitats or their ability to respond to environmental changes. To fill this knowledge gap, we conducted a litterbag decomposition experiment along a pronounced climate and vegetation gradient in the Chilean Coastal Cordillera (26°S to 38°S), ranging from hyper-arid to temperate, using mixtures of four plant species, native to the respective ecosystems. We analyzed potential decomposition functions of bacterial and fungal litter communities along with their biotic (litter traits) and abiotic (meteorological conditions and soil properties) environments, to determine the relative importance of these environmental factors for microbial community functioning. We also tested the impact of the functional diversity of the decomposer communities (i.e., the diversity of decomposition related functions) on litter mass loss.

Functional composition was related most strongly to the temporal variation of precipitation and radiation, explaining about 19 % and 6 % of variation in bacteria and fungi, respectively. In contrast, functional diversity was quite strongly related to litter chemical traits (C:N, C:P, tannins, phenols). Litter mass loss after six months of decomposition was not correlated to the functional diversity of decomposer communities but increased with the presence of habitat generalists like *Proteobacteria*, *Actinobacteria*, *Firmicutes*, and *Bacteroidetes*. Taken together, these results highlight i) the interplay between abiotic factors and chemical litter traits on litter microbial functions and functional diversity and ii) the importance of microbial generalists for litter decomposition across different ecosystems. These results enhance our ability to predict changes in microbial decomposer communities and litter decomposition under future climate-change scenarios.

## 1 Introduction

Plant litter decomposition is a crucial ecosystem process that links carbon (C) and nutrient cycling, impacting soil fertility and plant primary productivity (Aerts, 1997; Walter et al., 2020). This process is regulated by macro- and microclimate, litter traits (Canessa et al., 2021), as well as the abundance, diversity, structure, and activity of microbial decomposer communities (Bradford et al., 2017; Graham et al., 2016; Glassman et al., 2018; Waring et al., 2022). Previous work suggests that decomposition is controlled hierarchically by climate, substrate (i.e. litter traits), and microbes (Berg et al., 1993; Cornwell et al., 2008; Wang and Allison, 2022). However, these factors are not mutually independent and recent studies have challenged this order of importance, showing that microbial communities significantly influence decomposition patterns at regional scales (Bradford et al., 2016, 2017; Allison et al., 2013; Bray et al., 2012; Glassman et al., 2018; Gołębiewski et al., 2019; Martiny et al., 2017).

Functional traits of microbial communities have been shown to predict decomposition more accurately than microbial identity or community structure (Fanin and Bertrand, 2016; Huang et al., 2022; Schneider et al., 2012; Trivedi et al., 2016), because different taxonomic compositions do not necessarily reflect different functionalities (Keiser et al., 2011; Louca et al., 2018; Martiny et al., 2017; Schroeter et al., 2022). Taxonomic and functional diversity, however, are often related. High species diversity can lead to functional redundancy that allows communities to be resilient to environmental disturbances and maintain ecosystem functions, such as decomposition (Allison and Martiny, 2008; Shade et al., 2012). Still, approaches based on functional traits are often more informative than taxonomy-based approaches in assessing decomposition (Allison, 2012; Hu et al., 2019; Wang and Allison, 2022). For example, the diversity of the community-level catabolic ability may enhance decomposition in grasslands (Bonkowski and Roy, 2005) and the insight into microbial responses provided by analyzing microbial traits, such as functional genes, has been suggested to enhance our understanding of ecosystem functions (Hu et al., 2019; Shen et al., 2016). The impact of the functional traits of microbial decomposers and their diversity on litter decomposition rates, in interaction with climate and litter traits, however, is not well understood (Joly et al. 2023). This is unfortunate, considering that litter degradation substantially contributes to the terrestrial ecosystem’s CO_2_ emissions, a flux strongly challenging projections on the C cycle in Earth System models (Joly et al. 2023, Lennon et al. 2024).

Despite the recognition of the benefits of better understanding the functional traits of microbial decomposer communities, the drivers that influence microbial community functions and diversity in different ecosystems remain unclear. Variation imposed by environmental heterogeneity can shift microbial decomposer functions, with factors like moisture, temperature, land use, and soil properties explaining microbial functional variation across ecosystems (Chen et al., 2018; Elias et al., 2020; Fierer et al., 2012; Hu et al., 2019; Pailler et al., 2014; Paula et al., 2014; Rodriguez et al., 2022; Tringe et al., 2005; Wang et al., 2023). And not surprisingly, microbial functional diversity seems to be strongly related to plant community composition and litter trait diversity (Gillespie et al., 2021; Lange et al., 2015; Wang et al., 2019, 2020). For example, microbial communities exposed to poor quality litter likely exhibit a wider range of functions to utilize chemically diverse compounds than microbial communities exposed to high quality litter (van der Heijden et al., 2008; Keiser et al., 2011). Understanding the importance of these drivers for microbial community functions and diversity is crucial for predicting the community response to environmental change and for predicting litter decomposition patterns, as the drivers of microbial functional composition and diversity are ultimately the drivers of litter decomposition.

Functional diversity in microbial communities can be an indicator for their ability to adapt to environmental changes such as increasing temperature and drought, as higher functional diversity indicates a broad set of functions present in the community that allows it to cope with new conditions (Adje et al., 2023; Wallenstein and Hall, 2012). The presence and proportion of generalists and specialists can modulate community responses, with generalists better mitigating the impact of environmental change (Clavel et al., 2011; Wallenstein and Hall, 2012). Specialized decomposers can be highly efficient under the optimal conditions they developed on (Mouillot et al., 2011; Wallenstein and Hall, 2012), but generalist communities, on the other hand, can deal with various substrates and environmental conditions, important to maintain decomposition under unfavorable conditions (Bell and Bell, 2021; Osburn et al., 2022; Richmond et al., 2005).). The functional diversity of communities and their ability to maintain functions are important factors for predicting litter decomposition and nutrient dynamics under environmental change (Allison and Martiny, 2008; Fetzer et al., 2015; Osburn et al., 2022). The causal relationships, however, remain poorly understood restricting our capability for future projections of litter decomposition under changing environmental conditions. Therefore, while functional microbial community traits could significantly enhance biogeochemical models (Trivedi et al., 2013; Wieder et al., 2013), it is not yet possible to parameterize their impact on litter decomposition and element cycling quantitatively.

In this study, we investigate the impact of environmental parameters on microbial decomposer functions and diversity, and their role in litter decomposition. We did this by conducting a litterbag decomposition experiment along a climate and vegetation gradient in the Chilean Coastal Cordillera. The Chilean Coastal Cordillera provides a unique opportunity to study litter decomposition due to its wide spectrum of environmental conditions, ranging from arid deserts to temperate rainforests. We analyzed potential decomposition functions of bacterial and fungal litter communities, along with their biotic (litter traits) and abiotic environmental (meteorological conditions and soil properties) to investigate their relative importance for microbial communities. We asked the following question: How do the functional composition and diversity of bacteria and fungi in leaf litter interact with meteorological conditions and soil properties and correlate with litter decomposition rates along a long climate gradient in Chile? We hypothesized that 1) differences in chemical litter traits have a larger effect on microbial decomposition functions than meteorological conditions or soil properties, as specific chemical components may need specialized microorganisms to be broken down, that 2) differences in chemical litter traits affect microbial functional diversity more than abiotic conditions, and 3) litter mass loss would be positively correlated with functional diversity because a functionally diverse community would be able to efficiently decompose any litter, while a functionally uniform community would be able to efficiently decompose only certain compounds of the litter.

## 2 Methods

### 2.1 Study sites

The study was conducted along a continental transect in the Coastal Cordillera of Chile (26°S-38°S), covering diverse climates and ecosystems. Six sites were investigated with contrasting climate but similar granitoid parent material (Oeser et al., 2018). Detailed information about the vegetation, soil, and parent material at, or near, each study site can be found in Canessa et al. (2021, 2022), Übernickel et al. (2023), Bernhard et al. (2018), and Oeser et al. (2018). Briefly, the primary characteristics of each site are as follows (from north to south): Two sites were located in the western extend of the Atacama Desert within the National Park Pan de Azúcar (26°S, 71°W), including an arid-dry site (“AD”) in Pampa Blanca (little and irregularly influenced by fog), and a site in Las Lomitas, where incoming fog is an important resource of water (Lehnert et al., 2018), hence called “AF”, arid-fog. Moving south, the next sites are in a semi-arid shrubland (“SA”) in the Private Reserve Quebrada de Talca (30°S, 71°W), a mediterranean forest (“ME”) in the National Park La Campana (33°S, 71°W), an upland temperate forest (“TU”) in the National Park Nahuelbuta (38°S, 71°W) and in a lowland temperate rainforest (“TL”) in the Contulmo Natural Monument (38°S, 71°W). From desert to rainforests, the mean annual temperature decreases from 14.4°C to 6.7°C, while mean annual precipitation increases from 13 mm yr^-1^ to over 2 100 mm yr^-1^ (Canessa et al., 2022; van den Brink et al., 2023). The study sites were chosen to represent a wide range of climates and ecosystems, from the hyper-arid Atacama Desert to the temperate rainforests of southern Chile. Each site is characterized by its distinct climate, vegetation, and soil properties.

### 2.2 Experiment and sample collection

We conducted a litterbag decomposition experiment along the transect. Mixtures of four plant species (incl. one lichen in the case of AF), each native to and dominant in the respective site, were selected for the decomposition experiment and the subsequent analysis of microbial communities (Table 1). The 10×10 cm litter bags with 1 mm mesh size contained 2 g (±0.2 g) of oven-dried (at 60°C) litter, with each species contributing an equal amount. The bags were installed at the soil surface, which was cleared from litter, in early June 2017 in three independent plots per site. A detailed description of the species selection, the preparation of the litterbags and the installation can be found in Canessa et al. (2022). After 6 and 8 months of decomposition, i.e., late November/December 2017 and late January/February 2018, two litterbags were collected from each plot (74 litterbags in total). At each sampling plot and time, one litterbag was used for estimating litter mass loss (Canessa et al. 2022) and the other one was transferred to a nucleic acid preservation (NAP) buffer (Camacho-Sanchez et al., 2013; Menke et al., 2017) on-site and stored frozen upon arrival in the lab until the analysis.

**Table 1:** Selected species per site included in the litterbag decomposition ex-periment used for microbial analyses.

| Study site | Species |
| --- | --- |
| arid-dry desert (AD) | *Cristaria integerrima* Phil.<br>*Nolana crassulifolia* Poepp.<br>*Nolana mollis* (Phil.) Johnst.<br>*Tetragonia maritima* Barnéoud |
| arid-fog desert (AF) | *Leucostele deserticola* (Werderm.) Schlumpb.<br>*Euphorbia lactiflua* Phil.<br>*Nolana paradoxa* (Lindl.) Hook.<br>*Usnea eulychnae* |
| semi-arid shrubland (SA) | *Flourensia thurifera* DC.<br>*Gutierrezia resinosa* Blake<br>*Haplopappus decurrens* Rémy<br>*Senna cumingii* (Hook & Arn.) Irwin & Barneby |
| mediterranean forest (ME) | *Aristeguietia salvia* (Colla) King & Rob.<br>*Colliguaja odorifera* Molina<br>*Jubaea chilensis* (Molina) Baill.<br>*Quillaja saponaria* Molina |
| upland temperate rainforest (TU) | *Araucaria araucana* (Molina) Koch<br>*Gaultheria mucronata* (L.F.) Hook. & Arn.<br>*Nothofagus dombeyi* (Mirb.) Oerst.<br>*Nothofagus obliqua* (Mirb.) Oerst. |
| lowland temperate rainforest (TL) | *Aextoxicon punctatum* Ruiz & Pav.<br>*Greigia sphacelata* Regel<br>*Lophosoria quadripinnata* (Gmel.) C.Chr. in Skottsb.<br>*Nothofagus obliqua* (Mirb.) Oerst. |

### 2.3 Environmental parameters

#### Litter traits

Carbon, nitrogen, and phosphorus contents in litter, as well as the concentration of total phenols and tannins were determined by Canessa et al. (2021, 2022). A detailed description of the methods can be found in Canessa et al. (2021). Litter C:N and C:P ratios were calculated as measures of the substrate quality and total phenols and tannins were used as indicators of the substrate recalcitrance (Daebeler et al., 2022; Prescott, 2010; Talbot and Treseder, 2012). The contents and ratios of the four species per litterbag were used to calculate the means and the coefficients of variation (CV, Fig. S1) for each site that represent the trait variation within sites. In a few cases, if one species lacked a trait, the mean and CV were calculated with the remaining three species.

#### Chemical soil properties

Soil pH, soil C and N for the sites at the *arid-dry desert* and the *semi-arid shrubland* were determined in this study. Soil pH, soil C and N for the sites at the *mediterranean forest* and *upland temperate rainforest* were determined by Bernhard et al. (2018) in the upper 5 cm, and soil pH, soil C and N for the site at the *arid-fog desert* were determined by Jung et al. (2020) in the upper centimeter of the grit crusts covering the surface. Soil pH and soil C content from 0-35 cm depth in the soil for the *lowland temperate rainforest* were taken from Endlicher and Mäckel (1985). As no total N content was given by Endlicher and Mäckel (1985), the N content of the upland temperate rainforest was used to calculate soil C:N for the lowland temperate rainforest. Soil pH given as pH_H2O_ (AF, TL) was converted to pH_CaCl2_ by subtracting 0.5 (Davies, 1971).

#### Soil moisture and soil temperature

The volumetric soil moisture and soil temperature were recorded with TMS-4 datalogger (TOMST, Czech Republic) at each plot, directly next to the litterbags during the experiment (Canessa et al., 2022). Calibration for soil moisture was done using the Calibrator tool of the datalogger supplier using clay, silt and sand content obtained from Bernhard et al. (2018). We calculated the mean soil moisture and mean soil temperature as well as the coefficients of variation within each site for the periods 0-6 and 0-8 months as temporal variation within sites.

#### Meteorological parameters

Precipitation and solar radiation during the experiment period at the sites *arid-dry desert*, *semi-arid shrubland*, *mediterranean forest*, and *upland temperate rainforest* were obtained from the SPP-EarthShape (earthshape.net) weather stations (Übernickel et al., 2020). Precipitation data for the *arid-fog desert* (Las Lomitas) site was provided by Jörg Bendix (personal communication). For the *lowland temperate rainforest* (Contulmo), precipitation data was obtained from the Agrometerorología station La Isla, Purén (https://agrometeorologia.cl/) operated by the National Institute of Agricultural and Food Research and Technology (INIA). For the sites *arid-fog desert* and *lowland temperate rainforest*, the solar radiation from the *arid-dry desert* and *upland temperate rainforest*, respectively, was used. Sum of precipitation and mean solar radiation were calculated as well as their coefficients of variation for each site for the periods 0-6 and 0-8 months as temporal variation within sites.

### 2.4 Litter microbial community data

#### DNA extraction and sequencing

Total genomic DNA from litter material was extracted with the FastDNA^TM^ Spin Kit for Soil from MP Biomedicals (Santa Ana, California), following the manufacturer’s instructions. Prior to the extraction, the NAP was partially removed by centrifuging the material at 10000 rcf for 2 minutes and carefully withdrawing the supernatant/collected solution with a pipette. DNA purification, library preparation, and sequencing was done by the Institute for Medical Microbiology and Hygiene (MGM) at the University of Tübingen. The V4 region of the 16S rRNA gene and the ITS region were sequenced for the bacterial and fungal communities, respectively, using the MiSeq® Reagent Kit v3 (600 cycles) with on average 90k reads per sample.

#### Bioinformatic analyses

Raw data were processed with the nf-core/ampliseq pipeline (Ewels et al., 2020; Straub et al., 2020) by the Quantitative Biology Center (QBiC) Tübingen, including sequencing quality control with FastQC (Andrews, 2010), identification of amplicon sequence variants (ASVs) with DADA2 (Callahan et al., 2016), and the taxonomical classification of ASVs with QIIME2 (Bolyen et al., 2019). Non-target sequences, including chloroplasts and mitochondria, were removed from the analysis. Raw sequences were submitted to the NCBI database.

#### Prediction of decomposition functions

The prediction of functions related to litter decomposition was done by Tax4Fun2 (Wemheuer et al., 2020; Aßhauer et al., 2015) for bacterial communities, with Ref99NR as reference dataset. A subset of 71 functional orthologs were selected from the Kyoto Encyclopedia of Genes and Genomes (KEGG), representing enzymes known to degrade cellulose, hemicellulose, chitin, pectin, starch, or lignin (Table S1). The selection was based on Freedman et al. (2016). Additionally, 12 bacteria genera with known cellulolytic (and hemicellulolytic) capability (i.e., the capability to hydrolyze cellulose and/or hemicellulose; Table S2) were selected based on López-Mondéjar et al. (2016) and Tláskal et al. (2016). The prediction of ecological guilds in fungal communities was done via FUNGuild v 1.0 (Nguyen et al., 2015). For all statistical analyses, only the following guilds were included: Litter saprotrophs, leaf saprotrophs, wood saprotrophs, plant saprotrophs, soil saprotrophs, dung saprotrophs, and undefined saprotrophs. The abundance of taxa with multiple guild predictions was assigned equally to each guild.

#### Habitat generalists and specialists

Habitat generalists were identified as taxa with an incidence in more than 75% of the samples and a high relative abundance (>0.1%) over all sites (Rodriguez et al., 2022). Habitat specialists were defined by indicator-value analyses based on taxonomical annotated abundance tables, using *indval* from the package *labsdv* (Roberts and Roberts, 2016). Only taxa with a significant IndVal value > 0.7 were regarded as habitat specialists. Not all taxa fall into either category, i.e. part of the community is not represented by these categories.

### 2.5 Statistical analyses

All analyses were performed in R version 4.3.1 (R Core Team, 2023). The predicted decomposition functions of the microbial communities were compared between sites by non-metric multidimensional scaling (NMDS) based on Bray-Curtis distance matrices, using *metaMDS* from the R package *vegan* (Oksanen et al., 2007). Averaged subsampled Bray-Curtis dissimilarity matrices were calculated from the abundance of functions in the fungal communities using *avgdist* from *vegan*. Non-averaged Bray-Curtis dissimilarity matrices were calculated from relative abundances of functions within the bacterial community using *vegdist* from *vegan*. 2D stress of all NMDS was below 0.1. Differences between sites and months were tested by permutational multivariate analysis of variance using *adonis2* from *vegan* and a pairwise comparison of sites was done with *pairwise.adonis2* from the package *pairwiseAdonis* (Martinez Arbizu, 2017). The functional diversity of bacterial and fungal communities was evaluated based on the Shannon index. To test differences between sites and months we used ANOVA. Model assumptions (homogeneity of variance and normal distribution of residuals) and the presence of influential data points were checked visually with diagnostic plots. The abundance of cellulolytic genera in litter and in soil were square root transformed to meet the model assumptions. Differences between sampling times were not significant, which is why data from both time points were pooled for all further analyses.

#### Effect of environmental parameters

The variation of the predicted decomposition functions and the functional diversity between litter communities was partitioned between the environmental parameter sets *chemical litter traits* (i.e. C:N, C:P, tannins, phenols), *chemical soil properties* (soil C:N, soil pH), *thermal-moisture soil conditions* (soil moisture, soil temperature), and *meteorological conditions* (sum of precipitation, mean solar radiation). A distinction was made in separate partitioning between the between-site and within-site variation of the predictors. The between-site variation was represented by the site mean (respectively the sum of precipitation) and the within-site variation was represented by the coefficients of variation within each site. All explanatory variables were standardized prior to the analysis. The collinearity was checked based on variance inflation factors and soil temperature and soil pH (VIF>20) had to be excluded from the between-site model, to avoid saturation of the model. Partitioning was done with *varpart* from the package *vegan*. The fractions were tested with permutation tests, with 999 permutations. The fractions represent the adjusted percentage of either the unique contribution of one data set/predictor or the shared variation between two or more data sets. Following the same procedure, the variation in litter mass loss between sites was partitioned between the functional diversity and the percentage of habitat generalists and specialists (as well as abundance of cellulolytic bacteria genera) separately for each community. Additionally, Pearson’s correlation coefficients were calculated for the relation of the functional diversities, respectively litter mass loss, with the individual predictors of the significant environmental parameter sets.

## 3 Results

### 3.1 Decomposition functions and functional diversity

The bacterial litter communities showed a clear desert group, but, interestingly, the coolest and wettest site (upland rainforest) clustered in the middle of the axes, while site with a more intermediate climates (semi-arid and mediterranean) were located furthest from the desert group (Fig. 1a). Fungal litter communities (Fig. 1b) were clearly separated between a wet and a dry group, with the dry group including only the desert sites, based on their predicted decomposition functions.

**Figure 1:**
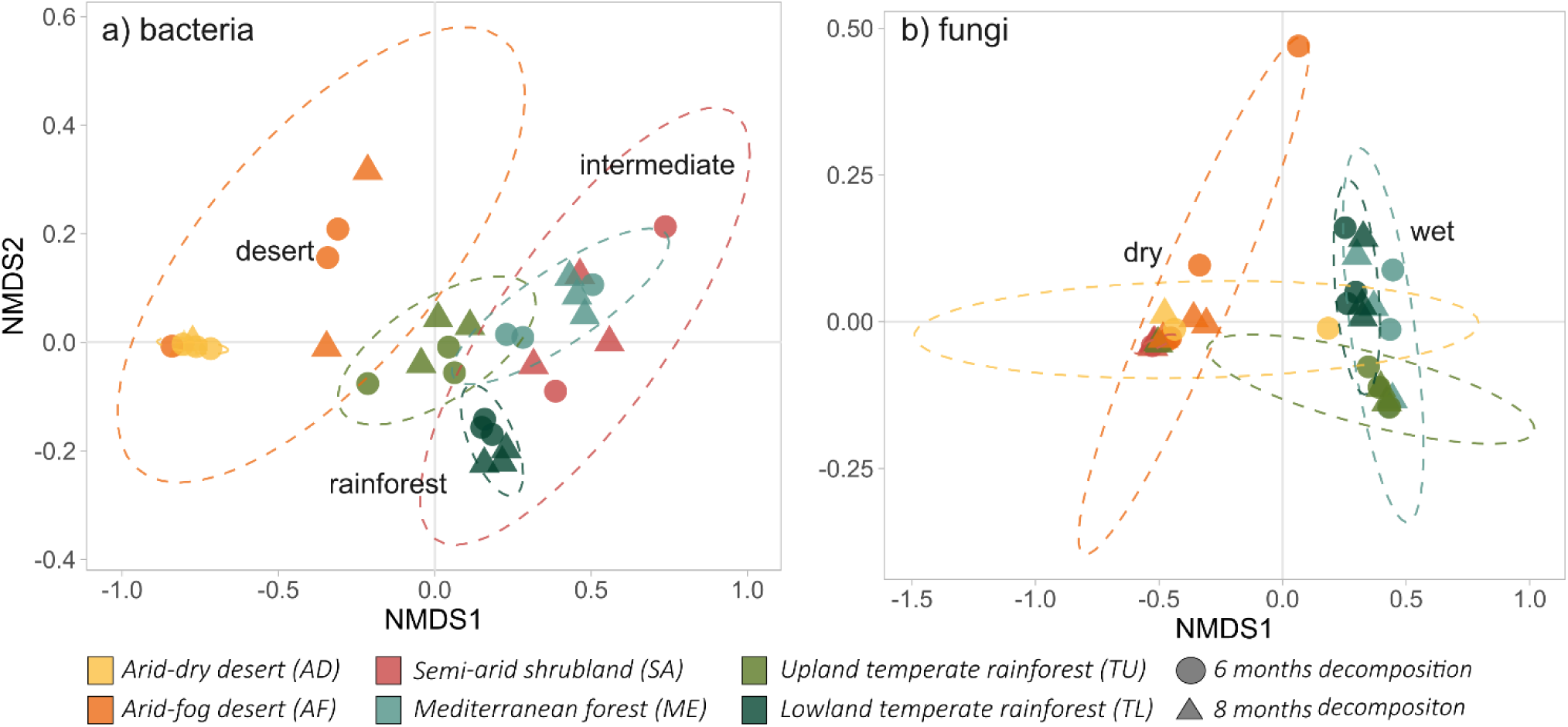
Functional composition of bacterial (a) and fungal (b) litter communities, displayed with non-metric multidimensional scaling (NMDS). NMDS for fungi are based on averaged Bray-Curtis dissimilarities (random subsampling). NMDS for bacteria are based on Bray-Curtis dissimilarities. Dashed lines = 95% confidence ellipses, assuming a multivariate t-distribution.

Along the climate gradient, bacterial functional diversity showed a unimodal pattern with a minimum at intermediate (i.e., semi-arid and mediterranean) conditions (Fig. 2a). The relative abundance of cellulolytic genera in bacterial litter communities (Fig. 2b) peaked in the mediterranean forest and was lowest in the arid dry site. Fungal functional diversity (Fig. 2c), in contrast, were similar across ecosystems, although the diversity at the lowland temperate rainforest differed from the upland temperate rainforest, the semi-arid shrubland, and the arid-dry desert.

**Figure 2:**
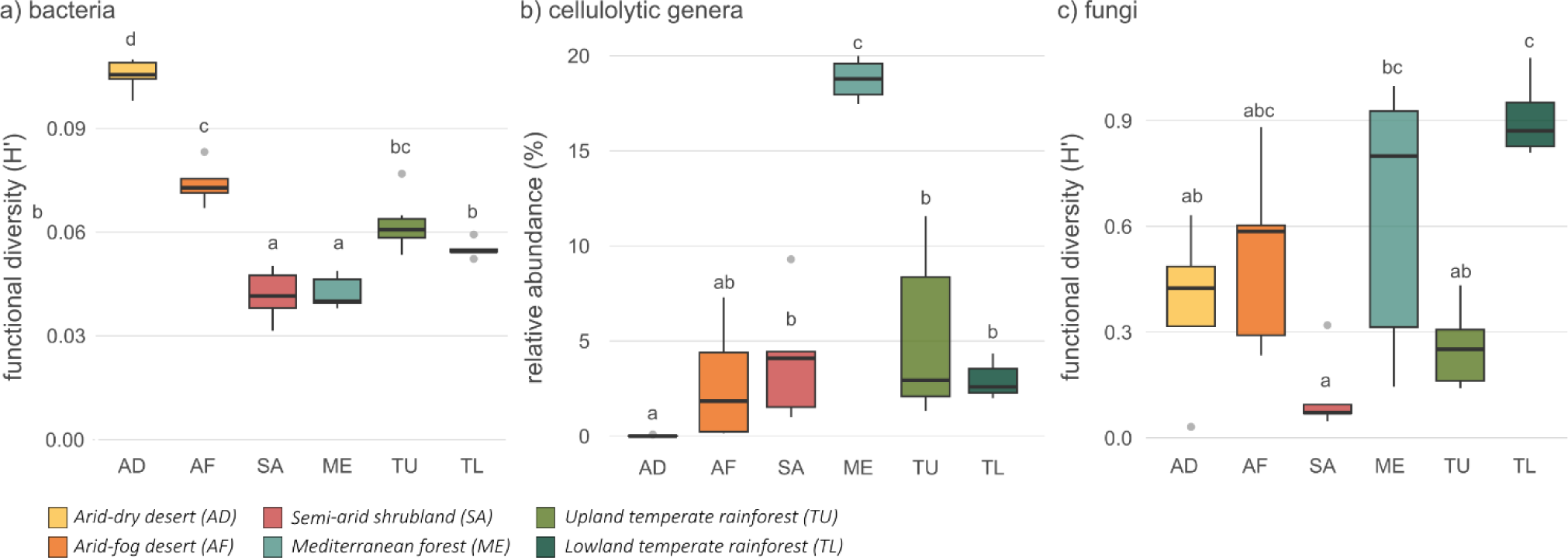
Functional diversity given as Shannon index (H’) of (a) bacterial litter communities and (c) of fungal litter communities. (b) Relative abundance of cellulolytic genera in bacterial litter communities. Different letters indicate significant differences between sites.

### 3.2 Habitat generalists and specialists

40 taxa among bacterial litter communities were identified as habitat generalists, and 11 taxa among fungal litter communities (Tab. S3). The driest site (AD) featured the highest percentage of bacterial (Fig. 3a) and fungal (Fig 3b) habitat generalists. From there, the percentage decreased towards the wettest site, the lowland temperate rainforest. At the arid-dry desert and the upland temperate rainforest, bacterial (Fig. 3a) and fungal (Fig. 3b) litter communities had the highest proportion of habitat specialists (i.e., indicator taxa). In absolute terms, most bacterial and fungal habitat specialists in litter communities were identified at the upland temperate rainforest (15 and 53, respectively), and none or close to none were detected among litter communities at the semi-arid shrubland, the mediterranean forest and the lowland temperate rainforest. A full list of the absolute numbers of habitat specialists and generalists, grouped by Phylum, is given in Table S3.

**Figure 3:**
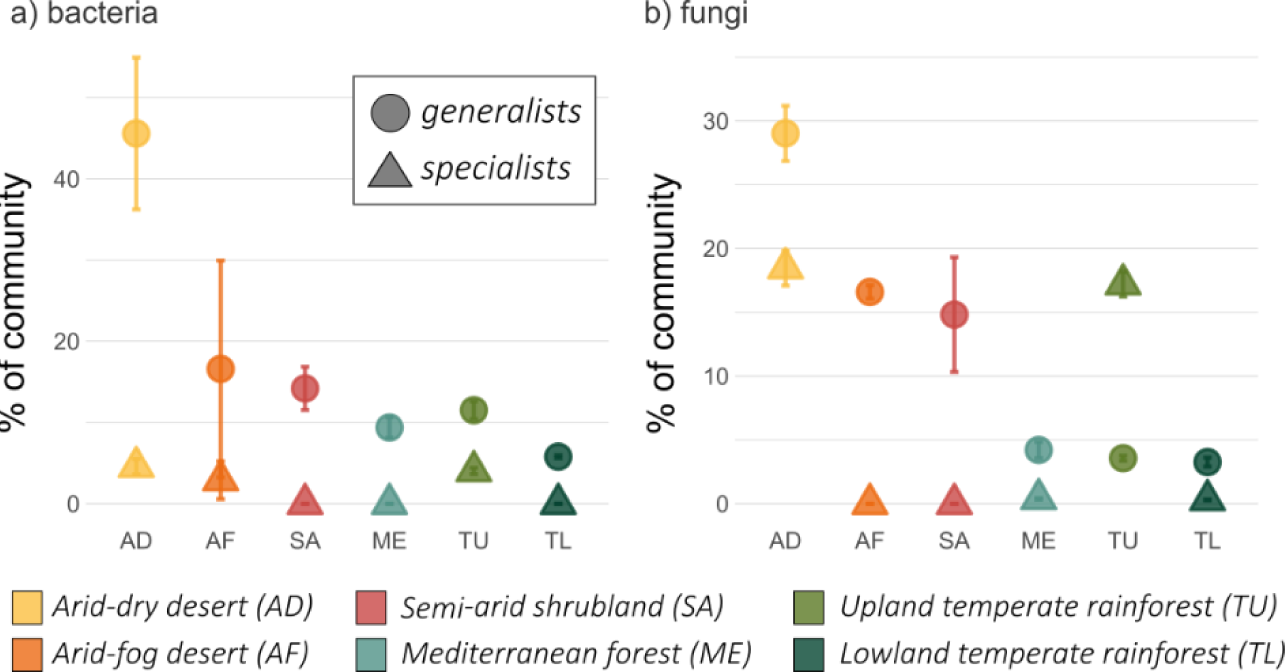
Percent of habitat generalists (circles) and habitat specialists IndVal; triangles) in bacterial (a) and fungal (b) litter communities. Taxa were identified as habitat generalists with an incidence of > 75% and a relative abundance of >0.1% over all samples. Taxa with an IndVal ≥ 0.7 were identified as indicator species = habitat specialists. Not all taxa fall into either category, e. part of the community is not represented by these categories.

### 3.3 Relative importance of environmental parameters in predicting decomposition functions and functional diversity

Environmental parameters, including chemical litter traits, chemical soil properties, thermal-moisture soil conditions, and meteorological conditions, were measured to understand their influence on microbial community decomposition functions. The temporal variation within each site of environmental conditions explained more of the variation in potential bacterial decomposition functions than between-site differences (Fig. 4a). However, differences in abiotic conditions between sites still explained more of the functional differences between bacterial communities than between fungal communities (Fig. 4b). And while all three between-site abiotic predictor sets explained differences between the bacterial communities, only chemical soil conditions (here soil C:N) explained differences between fungal communities. The temporal variation within each site of meteorological conditions (here precipitation and radiation) during the decomposition period was identified as the best predictor set for, both, bacterial and fungal communities. And while chemical litter trait variations explained 9% of the variation in potential decomposition functions between bacterial communities, none of the variation in potential fungal decomposition functions could be uniquely attributed to litter trait variations. The distance-based redundancy analysis (db-RDA) model that included all within-site predictor sets revealed that the within-site variation in radiation separated the bacterial desert communities, while the within-site variation of precipitation and soil moisture (and chemical litter traits) separated the mediterranean forest and semi-arid shrubland communities (Fig. S2c). Regarding the chemical litter traits, the variation of C:P ratios was most important, which was low for forest litter and high for litter of the arid-dry desert. The db-RDA model for the fungal communities showed the separation of the dry and wet cluster along a precipitation and radiation gradient (Fig. S2d).

**Figure 4:**
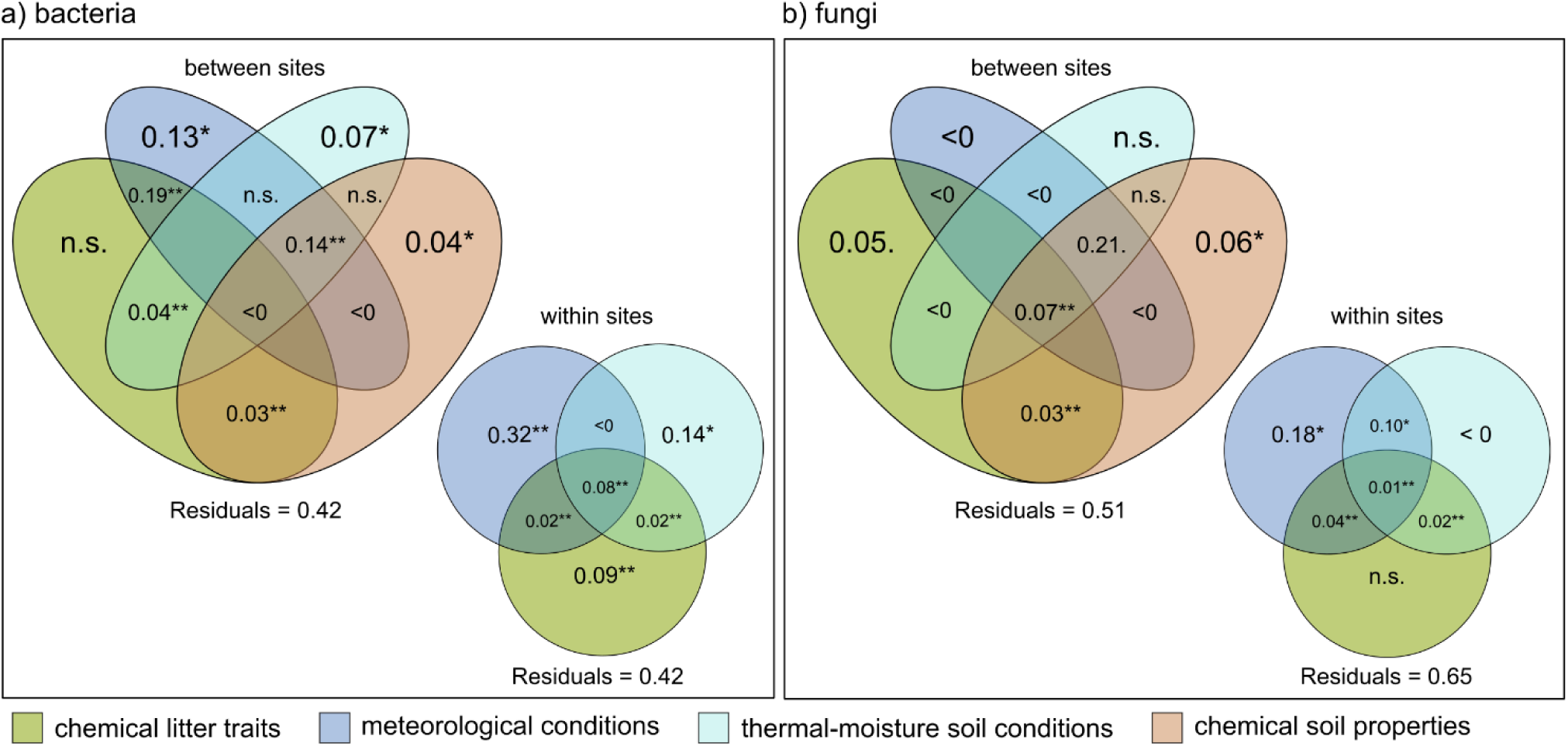
Partitioning of the variation of the microbial litter communities’ decomposition functions between the environmental explanatory data sets, separated into between-site (means) and within-site (CV) differences. Chemical litter traits were represented by C:N, C:P, tannins, and phenols, meteorological conditions were represented by precipitation and solar radiation, and thermos-moisture soil conditions were represented by soil moisture and soil temperature. Soil temperature and soil pH, however, had to be excluded from the between-site model due to collinearity, which saturated the model. Therefore, chemical soil properties are only represented by soil C:N and the thermal-moisture soil conditions are only represented by soil moisture. The total variation explained by the individual data sets can be subdivided into the fractions shown in the Venn diagram. The fractions represent the adjusted percentage of either the unique contribution of one data set or the shared variation between two or more data sets. The residual variation left unexplained is given below each figure. The fractions of contributions were tested with permutation tests with 999 permutations. Significance code: 0.001 ‘**’ 0.01 ‘*’ 0.05 ‘.’ 0.1 ‘ n.s.’ 1; ‘<0’ fraction contribution below zero; fractions not tested are not shown.

The unique contributions of the environmental parameters’ within-site variation to the functional diversity of bacterial communities were higher than the unique contributions of the parameters’ differences across sites (Fig. 5a). Differences of litter traits between sites and within sites explained the largest proportion of variation of bacterial as well as fungal functional diversity, respectively, followed by the temporal variation of precipitation and radiation during the decomposition period.

**Figure 5:**
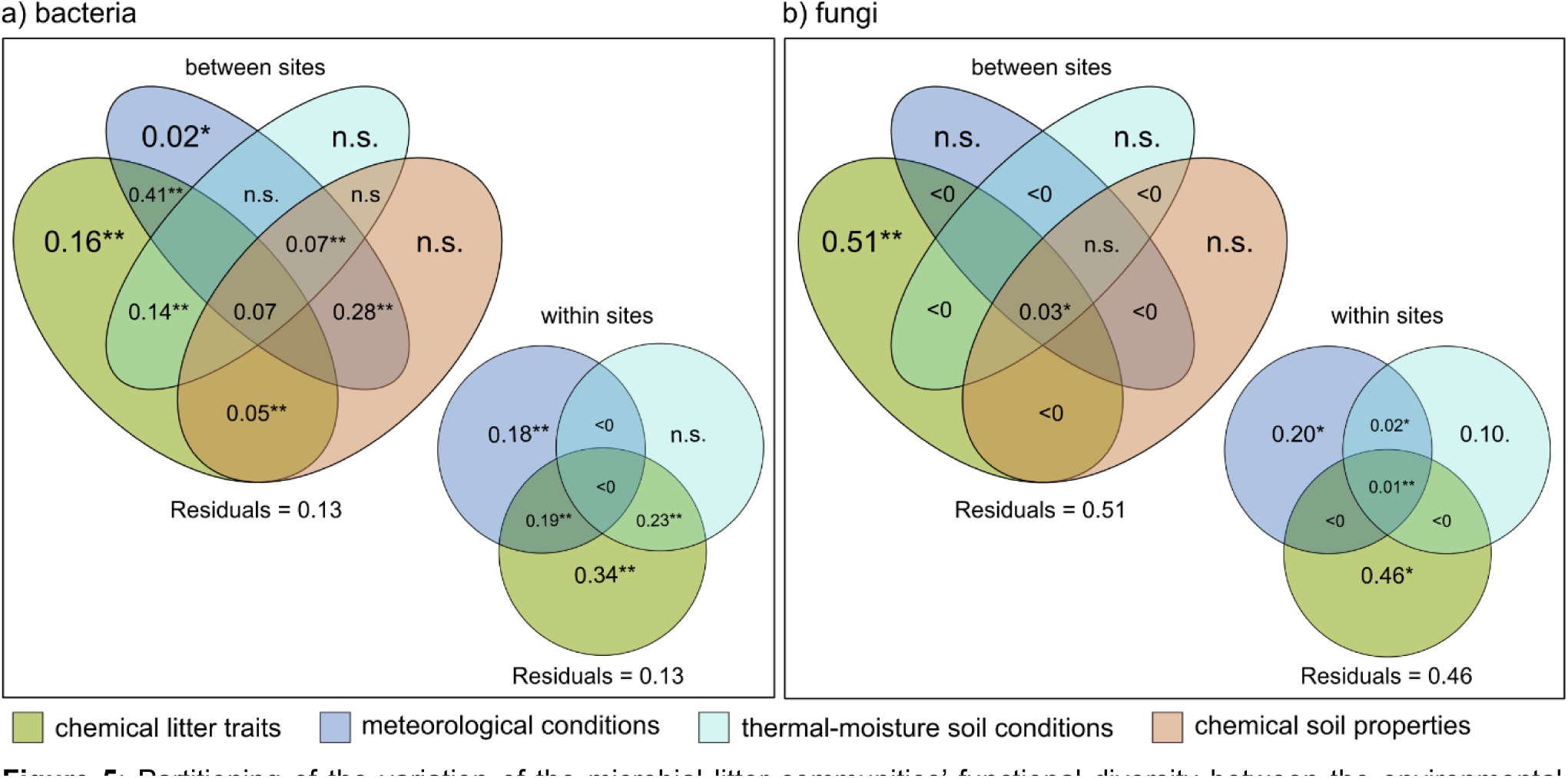
Partitioning of the variation of the microbial litter communities’ functional diversity between the environmental explanatory data sets, separated into differences across sites (means) and within sites (CV). Chemical litter traits were represented by C:N, C:P, tannins, and phenols, meteorological conditions were represented by precipitation and solar radiation, and thermos-moisture soil conditions were represented by soil moisture and soil temperature. Soil temperature and soil pH, however, had to be excluded from the between-site model due to collinearity. Therefore, chemical soil properties are only represented by soil C:N and the thermal-moisture soil conditions are only represented by soil moisture. The total variation explained by individual data sets can be subdivided into the fractions shown in the Venn diagram. The fractions represent the adjusted percentage of either the unique contribution of one data set or the shared variation between two or more data sets. The residual variation left unexplained is given below each figure. The fractions of contributions were tested with permutation tests. Significance code: 0.001 ‘**’ 0.01 ‘*’ 0.05 ‘.’ 0.1 ‘ n.s.’ 1; ‘<0’ fraction contribution below zero; fractions not tested are not shown

The functional diversity of bacteria negatively correlated with contents of tannins and phenols, while it correlated positively with the variation of C:N and tannins (Tab. 2). The functional diversity of fungi correlated positively with C:P ratios as well as the variation of phenol contents in litter. Large temporal variation of precipitation and soil moisture had a negative effect on the functional diversity of fungi and large temporal variation of radiation had a negative effect on the functional diversity of bacteria.

**Table 2:**
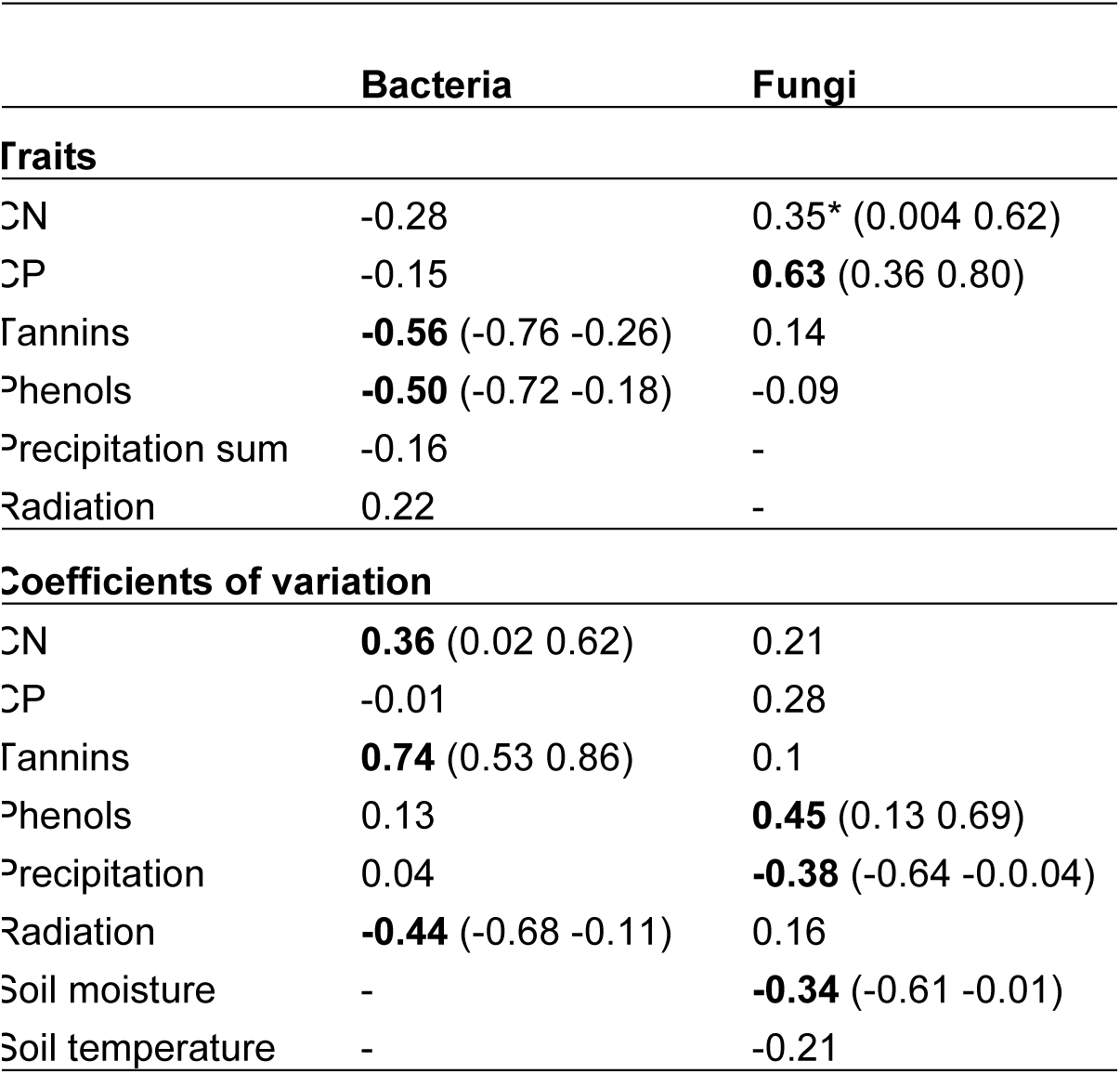
Correlation of the functional diversity of bacterial and fungal litter communities with means or sum (“traits”) and coefficient of variations per site of the environmental predictors. Pearson’s correlation coefficients are given. Significant correlations are indicated in bold, marginal significant correlations (p = 0.05) are indicated with asterisk, confidence intervals are given in brackets.

### 3.4 Relation of functions of microbial litter communities and litter mass loss

The litter mass loss of local plant species decomposing in their native habitats was highest in the arid-dry desert and the semi-arid forest (Fig. 6a). The percentage of generalists in bacterial and in fungal litter communities best explained differences of litter mass loss between sites, with a positive effect on mass loss (Tab. 3). Among the bacterial communities, the percentage of specialists had the second highest unique contribution. The functional diversity of, both, bacterial and fungal communities did not explain litter mass loss across sites.

**Figure 6:**
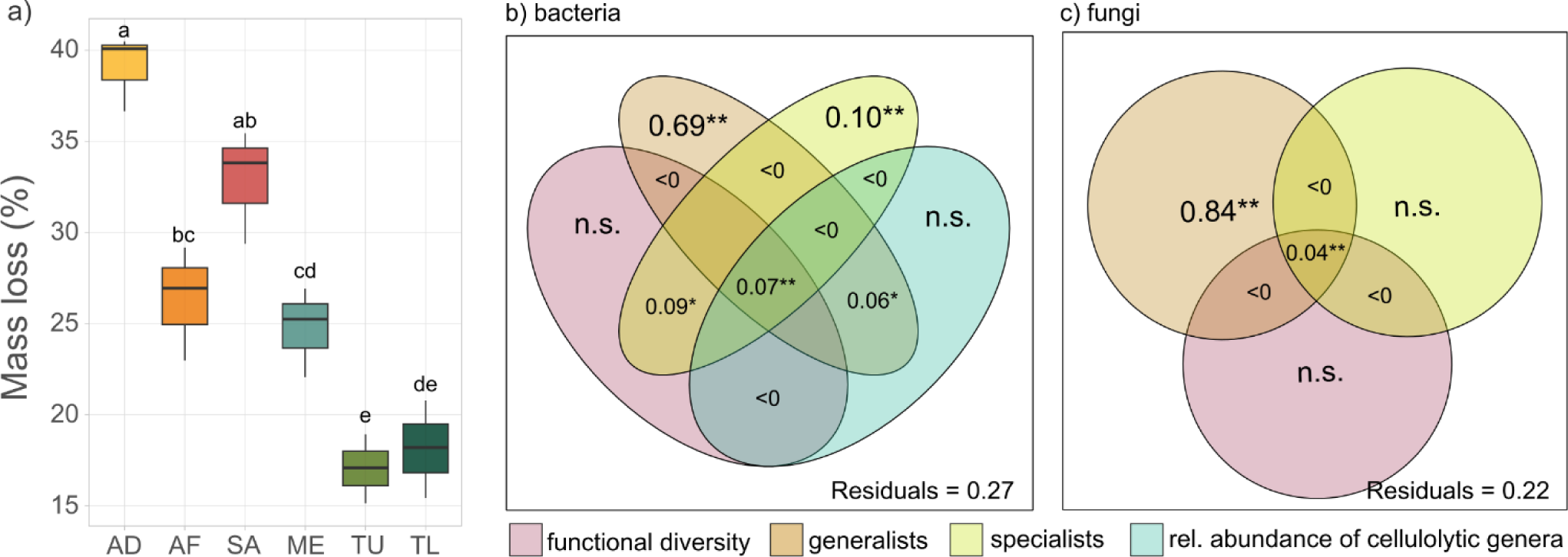
a) Mass loss per site after six months of decomposition determined in Canessa et al. (2022). Color code represent the different study sites, following figures 1-3. b+c) Partitioning of the variation in litter mass loss between characteristics of the litter microbial communities. The fractions represent the adjusted percentage of either the unique contribution of one predictor or the shared contribution of two or more predictors. The residual variation left unexplained is given in the lower righthand corner. The fractions of contributions were tested with permutation tests. Significance code: 0.001 ‘**’ 0.01 ‘*’ 0.05 ‘.’ 0.1 ‘ n.s.’ 1; ‘<0’ fraction contribution below zero; fractions not tested are not shown.

**Table 3:** Correlation of mass loss (%) with the proportion of generalists in the fungal and bacterial litter communities (%) and the proportion of specialists in the bacterial litter communities (%). Pearson correlation coefficients are given. Significant correlations are indicated in bold; confidence intervals are given in brackets.

|  | **Mass loss (%)** |
| --- | --- |
| bacterial generalists | **0.75** (0.43 0.90) |
| bacterial specialists | 0.20 |
| fungal generalists | **0.87** (0.69 0.95) |

## 4 Discussion

### 4.1 Effect of environmental parameters on potential decomposition functions

We hypothesized that differences in chemical litter traits across study sites would have a greater effect on the composition of potential decomposition functions of microbial litter communities than meteorological conditions or soil properties (*hypothesis 1*). However, differences in chemical litter traits between sites did not uniquely explain the variation in bacterial decomposition functions. Instead, within-site litter differences emerged as significant large-scale determinants, albeit to a lesser extent than abiotic factors. Our results suggest that between-site differences and temporal variations in abiotic habitat conditions shape the composition of bacterial functions stronger than litter traits. We further observed that chemical soil properties dominantly shaped the composition of fungal decomposition functions. This is in line with observations on bacterial litter communities that expressed similar catabolic structures across different litter types but varied with precipitation changes (Kheir et al., 2021) and with studies on soil microbial functions that changed with edaphic conditions (Hu et al., 2019; Wang et al., 2023; Yang et al., 2014) and micro(climate) (Chen et al., 2018; Hartmann et al., 2013). This suggests climatic conditions as important predictor to be considered for projections of litter decomposition in space and time. However, contrary to our findings, litter trait variation was found to decrease functional genes for carbohydrate and amino acid metabolism in bacterial litter communities and increased them in fungal litter communities, indicating significant litter trait effects on the functional potential of litter decomposer communities (Wang et al., 2019, 2020).

The functional composition of fungi related to the temporal variations in habitat conditions less than the functional composition of bacteria, suggesting fungal litter communities to be either less immediately impacted by large-scale abiotic constraints or to respond slower than bacterial communities to new conditions. Literature on community composition point to a higher taxonomic responsiveness of fungal litter communities than bacterial litter communities to environmental changes such as vegetation succession (e.g., Zhong et al., 2018; Zhan et al., 2021) or increased soil temperature and altered precipitation (Wahdan et al., 2022). Likewise, Martiny et al. (2017) found grassland litter bacterial composition to be more resilient than fungal composition to environmental change, such as precipitation reductions. However, bacterial functioning was not resilient (Martiny et al., 2017). Drought increased the abundance of genes targeting starch, xylan, and mixed polysaccharides in the bacterial community, while decreasing the abundance of genes targeting oligosaccharides, leading to lower decomposition rates in response to drought (Martiny et al., 2017). Thus, the stronger impact on the functional composition of bacteria in our study can indicate a faster functional adaptation to new environmental conditions.

Our findings indicate that conclusions drawn from drivers of taxonomic community composition or abundance cannot easily be transferred to the composition of potential decomposition functions of communities (see also e.g., Castro et al., 2009; Purahong et al., 2014, Shen et al., 2016; Yue et al., 2015) and provide further evidence of distinct responses of fungi and bacteria in their potential to decompose litter under environmental changes, with bacteria and their functions related to litter decomposition showing a higher response to changing (micro-)climatic conditions than fungi. This indicates that climate-driven changes in litter decomposition may mainly originate from altered bacterial communities and their functions, calling for a focused monitoring of bacterial litter decomposition traits along space-for-time chronosequences but also in climate-manipulating long-term experiments on litter decomposition.

### 4.2 Effect of environmental parameters on functional diversity

In contrast to functional composition, the differences of litter traits (C:N, C:P, tannins, phenols) within and between sites explained the largest proportion of functional diversity in bacterial and fungal litter communities. Regional variations in precipitation and radiation had a significant, though lesser, impact on functional diversity. This confirms our hypothesis that within-site variation of chemical litter traits affects microbial functional diversity more than abiotic environmental conditions (*hypothesis 2*). This aligns with earlier findings that litter traits had a stronger effect on litter and soil microbial functional diversity than edaphic factors (Shen et al., 2016; Xiao et al., 2018). As the dominant determinant across study sites, litter characteristics play a crucial role in determining microbial functional diversity on a large scale. Functional diversity of bacterial and fungal communities was lowest in the intermediate sites, distinguishing these intermediate sites from the gradient’s endmembers. In the extreme desert sites, contrary to traditional environmental filtering theory (e.g., Weiher et al., 1995), high litter diversity was found, such as succulent leaves and cactus needles (Canessa et al., 2022). Similarly, temperate rainforests exhibited high litter diversity with easy-to-decompose deciduous leaves and tough conifer needles. This diverse litter input likely promotes higher functional diversity in desert and rainforest sites, as microbial communities adapt to utilize chemically diverse compounds (van der Heijden et al., 2008; Keiser et al., 2011; Keiser and Bradford, 2017). Litter quality, however, had differential effects on the functional diversity of bacteria and fungi. While low litter quality (with high tannin and phenol contents) decreased functional diversity of bacteria, low litter quality with high C:P ratios increase the functional diversity of fungi. Further, large differences in litter C:N and content of tannins, increased functional diversity of bacteria, while large differences in contents of phenols increased the functional diversity of fungi.

### 4.3 Impact of on microbial decomposer functions and diversity on litter mass loss across ecosystems

We hypothesized that litter mass loss would correlate positively with functional diversity (*hypothesis 3*). However, litter mass loss could not be uniquely attributed to the functional diversity of decomposer. This unresponsiveness may be due to functional redundancy or differential stimulation of microbial decomposers by abiotic factors across sites (Pioli et al., 2020). Keiser and Bradford (2017) found that soil microbial functions predicted litter mass loss only in later decomposition stages. Thus, our study period of six months might have been too short to capture the potential of the community functional diversity for litter decomposition. Further, the ability of the decompose community is negatively correlated with litter quality has to be considered. Litter of high quality can be decomposed by communities with low ability (van den Brink et al., 2023), but litter mass loss is still fast as the litter is easy to decompose. Litter of low quality, as in the desert and rainforest, needs specialists with a high decomposer ability – and although the functional diversity is then higher, decomposition will be slower due to the low quality of the litter (van den Brink et al., 2023). Functional diversity, therefore, can be considered to be the result of litter quality (Fig. 5) but it cannot be distinguished as large-scale driver of litter mass loss by our study design. Habitat specialists and habitat generalists were better predictors of litter mass loss than the functional diversity. Habitat specialists in bacterial communities do affect litter mass loss positive. However, it is the proportion of habitat generalists in both bacterial and fungal communities that had the strongest positive impact.

The generalists, present in high abundance across all litterbags, were dominated by *Proteobacteria* and *Actinobacteria*, followed by *Firmicutes* and *Bacteroidetes*. This aligns with findings that *Proteobacteria*, *Actinobacteria*, and *Bacteroidetes* are the dominant bacteria phyla on decomposing litter (Purahong et al. 2016; Tlaskal et al., 2016; Urbanová et al., 2015). Genera from all four phyla produce extracellular enzymes for active substrate degradation (Buresova et al., 2019) and *Proteobacteria, Actinobacteria* and *Firmicutes* are known to be able to degrade recalcitrant compounds such as lignin (Bugg et al., 2018; Buresova et al., 2019; Větrovský et al., 2014; Zimmermann, 1990). *Proteobacteria* were shown to dominate in the early decomposition phase giving way to *Actinobacteria* (Schroeter et al., 2022). Other studies showed that *Bacteroidetes*, known for their decomposing activity, dominate in the first three months of litter decomposition, followed by *Proteobacteria* in the mid-stage (Buresova et al., 2019), which is in line with our findings after six months of decomposition. *Proteobacteria* and *Actinobacteria* were also shown to dominant irrespective of litter type, which identified them as early and secondary generalist decomposers, respectively (Buresova et al., 2019; Schroeter et al., 2022; Šnajdr et al. 2011). Fungal generalists were mainly genera from *Ascomycota* and *Basidiomycota,* known to be abundant decomposer able to utilize various carbon sources (Lopez-Mondejar et al., 2018). Bacterial and fungal generalists increased litter mass loss indicating their importance for the early to mid-decomposition phase across ecosystems and their importance in maintaining decomposition as an ecosystem function.

### 4.4 Implications

Ecosystem stability refers to the capacity of an ecosystem to withstand disturbances or return to balance afterward (Shade et al., 2012). Microbial community functioning plays a crucial role in maintaining this stability in the face of anthropogenic and environmental stress (Osburn et al., 2023). Given the dominant influence of precipitation and radiation on bacterial functional composition, the predicted decrease in precipitation of up to 50% until 2080-2100 (Christensen et al., 2007; Garreaud 2011) will impact ecosystem functioning both directly through altered water availability and indirectly by affecting bacterial litter communities. Functional diversity serves as an indicator of litter communities’ responses to environmental changes such as increasing temperatures and drought. Higher functional diversity enhances the ability to cope with or adapt to new conditions (Adje et al., 2023). Our results suggest that microbial decomposer communities will respond most strongly to shifts in plant community composition (i.e., changes in chemical litter traits) with regard to their functional diversity, rather than directly to meteorological changes. Shifts in plant species diversity significantly impact litter microbial communities (Santonja et al., 2017, 2018), suggesting that predicted climatic shifts will have stronger indirect effects through changes in litter composition than direct effects.

Different microbial taxa are responsible for different ecosystem functions (Wagg et al., 2019), emphasizing the importance of functional diversity. Habitat generalists, capable of tolerating a broad range of environmental conditions and utilizing diverse substrates, are crucial for sustaining ecosystem functions under adverse conditions (Bell and Bell, 2021; Wallenstein and Hall, 2012). Therefore, litter communities with higher functional diversity and a greater proportion of habitat generalists, as observed at the desert sites, may be more resilient to change and nutrient and carbon cycling in these systems is likely less impacted.

While functional breath of soil microbial decomposer communities was shown to be important for predicting litter decomposition and nutrient dynamics under environmental change (Osburn et al., 2022), we found no direct correlation between litter mass loss and the functional diversity of bacterial or fungal litter communities. The decomposer functional diversity may be crucial for maintaining decomposition as the environment changes onsite but might not necessarily be a direct predictor of decomposition rates on larger scales.

Shifts in fungal to bacteria ratios have been associated with both higher (Wardle et al. 2004) and lower (Ramirez et al. 2012) rates of litter decomposition. When bacterial and fungal litter communities do not only respond in abundance but also in their ability to decompose litter differently to abiotic environmental changes, this will impact litter decomposition further (Fischer et al., 2006). If bacterial litter communities respond and adapt more rapidly to changes, and thereby are able to maintain decomposition while fungi lag behind, the impact of environmental changes on carbon and nutrient cycling may be low. However, shifts in their respective functions will also affect their evolved interactions (Wagg et al., 2019; Wahdan et al., 2022) and a misaligned shift in the decomposition abilities of bacteria and fungi might hamper the overall decomposition process.

## 5 Conclusion

Our findings indicate that abiotic factors dominate microbial functional composition, but that substrate dominate microbial functional diversity. Our findings indicate that conclusions drawn from drivers of community composition or abundance cannot easily be transferred to potential functions of decomposer communities and provide further evidence of distinct responses of fungi and bacteria not only in composition and abundance but also in their potential ability to decompose litter under environmental changes, with a supposed higher sensitivity of bacterial functions to climate change. Bacterial and fungal generalists known for their role in litter decomposition were better predictors for litter decomposition in the first six months than microbial functional diversity. This enhances our ability to predict changes in litter decomposition under climate change, as microbial functional diversity and generalist presence are crucial indicators of ecosystem stability and resilience.

## Acknowledgements

We thank the Chilean National Park Service Corporacion Nacional Forestal (CONAF) for the permission to work in the National Parks Pan de Azúcar, La Campana, and Nahuelbuta. We also thank the agricultural community Quebrada de Talca and the rangers from the national parks for access to the study sites, and assistance in the field. This study was funded by the German Research Foundation (DFG) Priority Program SPP-1803 “EarthShape: Earth Surface Shaping by Biota” (BA 3843/6-1, KU 1184/36-1, DI 2136-11, NE 1852/3-2, OE 516/7-1 and -2, TI 338/14-1 and -2, SCHO 739/17). We further thank the excellence strategy of the University Tübingen for funding the work of S.C. Stock. L. van den Brink thanks additional support from ANID Anillo Act210038 funding the Institute of Ecology and Biodiversity (Chile). We thank the Quantitative Biology Center (QBiC), Institute for Medical Genetics and Applied Genomics (IMGAG) and Institute for Medical Microbiology and Hygiene (MGM) at the University of Tübingen for supporting the DNA amplicon sequencing and the primary bioinformatical data processing.

