## Supplement for "Environmental versus litter traits as drivers of microbial decomposer functions"

### Supporting information

Table S1: List of the selected functional orthologs from the Kyoto Encyclopedia of Genes and Genomes (KEGG), representing enzymes known to degrade cellulose, hemicellulose, chitin, pectin, starch, or lignin. The prediction of functions related to litter decomposition was done by Tax4Fun2 (Wemheuer et al., 2020) for bacterial communities, with Ref99NR as reference dataset. The selection was based on Freedman et al. (2016).

| Substrate | Functional ortholog | Enzyme | EC Number |
|---|---|---|---|
| Cellulose | K05349 | Beta-glucosidase | 3.2.1.21 |
| | K01188 | Beta-glucosidase | 3.2.1.21 |
| | K05350 | Beta-glucosidase | 3.2.1.21 |
| | K01179 | endoglucanase | 3.2.1.4 |
| | K19357 | cellulase | 3.2.1.4 |
| | K20542 | endoglucanase | 3.2.1.4 |
| | K01225 | Cellulose 1,4-beta-cellobiosidase | 3.2.1.91 |
| | K19668 | Cellulose 1,4-beta-cellobiosidase | 3.2.1.91 |
| | K20829 | Cellulose 1,4-beta-cellobiosidase | 3.2.1.176 |
| | K01180 | endo-1,3(4)-beta-glucanase | 3.2.1.6 |
| | K19069 | cellobiose oxidase | 1.1.99.18 |
| | K00702 | cellobiose phosphorylase | 2.4.1.20 |
| Hemicellulose | K01189 | alpha-galactosidase | 3.2.1.22 |
| | K07406 | alpha-galactosidase | 3.2.1.22 |
| | K07407 | alpha-galactosidase | 3.2.1.22 |
| | K01235 | alpha-glucuronidase | 3.2.1.139 |
| | K01209 | alpha-L-arabinofuranosidase | 3.2.1.55 |
| | K15921 | arabinoxylan arabinofuranohydrolase | 3.2.1.55 |
| | K20844 | non-reducing end alpha-L-arabino-furanosidase | 3.2.1.55 |
| | K01190 | beta-galactosidase | 3.2.1.23 |
| | K12111 | evolved beta-galactosidase subunit alpha | 3.2.1.23 |
| | K12308 | beta-galactosidase | 3.2.1.23 |
| | K12309 | beta-galactosidase | 3.2.1.23 |
| | K01192 | beta-mannosidase | 3.2.1.25 |
| | K01198 | xylan 1,4-beta-xylosidase | 3.2.1.37 |
| | K15920 | xylan 1,4-beta-xylosidase | 3.2.1.37 |
| | K22268 | xylan 1,4-beta-xylosidase | 3.2.1.37 |
| | K01181 | endo-1,4-beta-xylanase | 3.2.1.8 |
| | K13465 | ethylene-1,4-beta-xylanase | 3.2.1.8 |
| | K09252 | feruloyl esterase | 3.1.1.73 |
| | K21016 | feruloyl esterase | 3.1.1.73 |
| | K01218 | mannan endo-1,4-beta-mannosidase | 3.2.1.78 |
| | K19355 | mannan endo-1,4-beta-mannosidase | 3.2.1.78 |
| | K20832 | galactan endo-beta-1,3-galactanase | 3.2.1.181 |
| | K06113 | arabinan endo-1,5-alpha-L-arabinosidase | 3.2.1.99 |
| | K01224 | arabinogalactan endo-1,4-beta-galacto-sidase | 3.2.1.89 |

| | | | |
|---|---|---|---|
| | K20845 | endo-1,3-beta-xylanase | 3.2.1.32 |
| | K15531 | oligosaccharide reducing-end xylanase | 3.2.1.156 |
| | K18576 | xyloglucan endo-beta-1,4-glucanase | 3.2.1.151 |
| | K18578 | xyloglucan-specific exo-beta-1,4-glucanase | 3.2.1.155 |
| Chitin | K01205 | Alpha-N-acetylglucosaminidase | 3.2.1.50 |
| | K01452 | Chitin deacetylase | 3.5.1.41 |
| | K01183 | chitinase | 3.2.1.14 |
| | K13381 | bifunctional chitinase/lysozyme | 3.2.1.14 |
| | K20547 | basic endochitinase B | 3.2.1.14 |
| | K01207 | beta-N-acetylhexosaminidase | 3.2.1.52 |
| | K12373 | hexosaminidase | 3.2.1.52 |
| | K14459 | hexosaminidase | 3.2.1.52 |
| | K22278 | peptidoglycan-N-acetylglucosamine deacetylase | 3.5.1.104 |
| | K01567 | peptidoglycan-N-acetylmuramic acid deacetylase | 3.5.1.- |
| Pectin | K01184 | polygalacturonase | 3.2.1.15 |
| | K01213 | galacturan 1,4-alpha-galacturonidase | 3.2.1.67 |
| | K22933 | galacturan 1,4-alpha-galacturonidase | 3.2.1.67 |
| | K01728 | pectate lyase | 4.2.2.2 |
| | K22539 | pectate lyase | 4.2.2.2 |
| | K01732 | pectin lyase | 4.2.2.10 |
| | K01051 | pectinesterase | 3.1.1.11 |
| Starch | K01176 | alpha-amylase | 3.2.1.1 |
| | K05343 | maltose alpha-D-glucosyltransferase / alpha-amylase | 3.2.1.1 |
| | K07405 | alpha-amylase | 3.2.1.1 |
| | K01187 | alpha-glucosidase | 3.2.1.20 |
| | K12316 | lysosomal alpha-glucosidase | 3.2.1.20 |
| | K12317 | neutral alpha-glucosidase C | 3.2.1.20 |
| | K01178 | glucoamylase | 3.2.1.3 |
| | K21574 | glucan 1,4-alpha-glucosidase | 3.2.1.3 |
| Lignin | K00421 | laccase | 1.10.3.2 |
| | K05909 | laccase | 1.10.3.2 |
| | K23515 | lignin peroxidase | 1.11.1.14 |
| | K20205 | manganese peroxidase | 1.11.1.13 |

Table S2: Selected bacterial genera with known cellulolytic capability based on López-Mondéjar et al. (2016) and Tláskal et al. (2016).

| **Phylum** | **Genus** |
|---|---|
| *Proteobacteria* | *Dyella* |
| *Proteobacteria* | *Erwinia* |
| *Proteobacteria* | *Frateuria* |
| *Actinobacteriota* | *Frigoribacterium* |
| *Proteobacteria* | *Luteibacter* |
| *Bacteriodota* | *Mucilaginibacter* |
| *Firmicutes* | *Paenibacillus* |
| *Proteobacteria* | *Paucibacter* |
| *Bacteriodota* | *Pedobacter* |
| *Proteobacteria* | *Pseudoxanthomonas* |
| *Bacteriodota* | *Solitalea* |
| *Bacteriodota* | *Sphingobacterium* |

Table S3: Indicator (IndVal) and generalist analysis for taxa of bacterial (a) and fungal (b) litter and soil communities. Species with an IndVal >= 0.7 were identified as indicator species (i.e., habitat specialists). Species were identified as generalists with an incidence of > 75% and a relative abundance of >0.1% over all samples (i.e., habitat generalists). Numbers of taxa per Phylum.

| | ***Specialists*** | | | | | | | | | | | | ***Generalists*** | |
|---|---|---|---|---|---|---|---|---|---|---|---|---|---|---|
| | *Litter* | | | | | | *Soil* | | | | | | *Litter* | *Soil* |
| | *AD* | *AF* | *SA* | *ME* | *TU* | *TL* | *AD* | *AF* | *SA* | *ME* | *TU* | *TL* | | |
| | | | | | | | ***Bacteria*** | | | | | | | |
| *Acidobacteriota* | | | | | | | | | | | | 3 | | 6 |
| *Actinobacteria* | | 2 | | | 3 | | 1 | | | | | | 10 | 36 |
| *Armatimonadota* | | | | | | | | | | | | | | 1 |
| *Bacteroidota* | | 1 | | | | | | | | | 2 | 1 | 5 | 12 |
| *Bdellovibrionota* | | | | | | | | | | | | | | 1 |
| *Chloroflexi* | 1 | | | | | | | | | | | 2 | | 10 |
| *Crenarchaeota* | | | | | | | | | | | | | | 1 |
| *Elusimicrobiota* | | | | | | | | | | | | 1 | | |
| *Firmicutes* | 2 | | | | | | | | | | | | 7 | 5 |
| *Gemmatimonadota* | | | | | | | | | | 1 | | | | 3 |
| *Myxococcota* | | | | | | | | | | | | 1 | | 6 |
| *Patescibacteria* | | | | | | | | | | | | 1 | | 2 |
| *Planctomycetota* | | | | | | | | | | | | 1 | | 8 |
| *Proteobacteria* | 1 | 4 | | | 11 | | | 2 | | | 4 | 3 | 18 | 32 |
| *Spirochaetota* | | | | | | | | | | | | 2 | | |
| *Verrucomicrobiota* | | | | | 1 | | | | | | 1 | 2 | | 4 |
| ***Total*** | 4 | 7 | 0 | 0 | 15 | 0 | 1 | 2 | 0 | 1 | 7 | 17 | 40 | 127 |
| | | | | | | | ***Fungi*** | | | | | | | |
| *Ascomycota* | 6 | | | | 28 | 1 | 5 | 8 | 1 | | 6 | | 9 | 7 |
| *Basidiomycota* | | | | | 23 | | | | | | 1 | | 2 | 1 |
| *Chytridiomycota* | | | | 1 | 2 | | | | | | 1 | | | |
| *Mortierellomycota* | | | | | | | | | | | | | | 1 |
| *Olpidopmycota* | 1 | | | | | | | | | | | | | |
| ***Total*** | 7 | 0 | 0 | 1 | 53 | 1 | 5 | 8 | 1 | 0 | 8 | 0 | 11 | 9 |

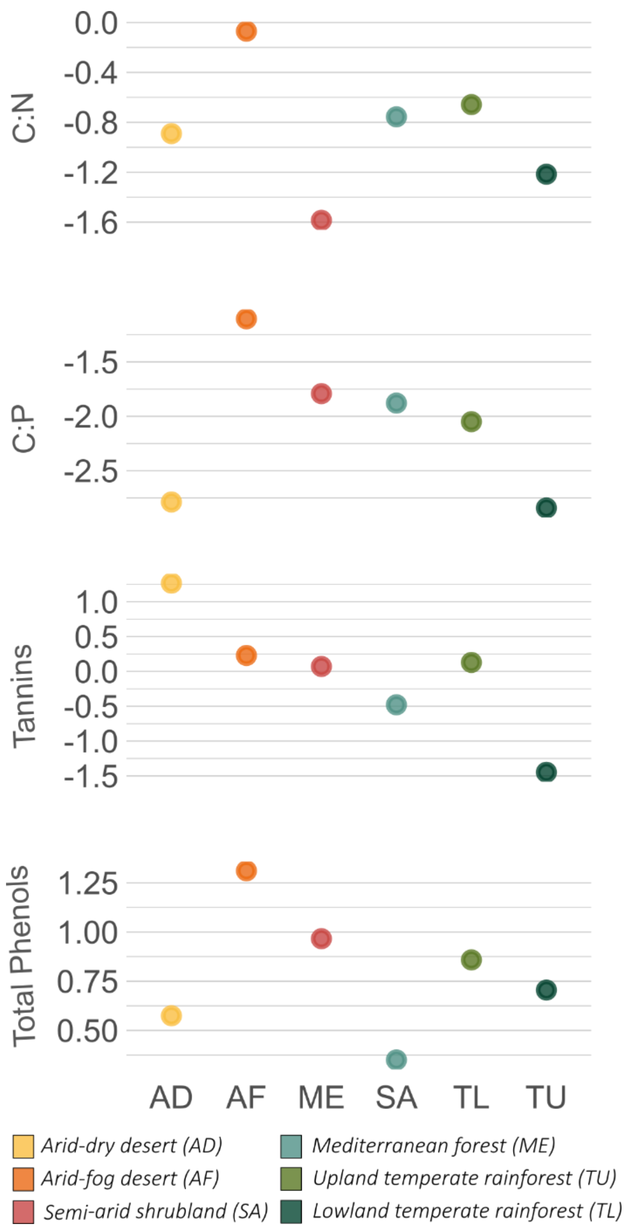


Figure S1: Overview of litter traits. CVs from four plant species used in the litterbag decomposition experiment per site.

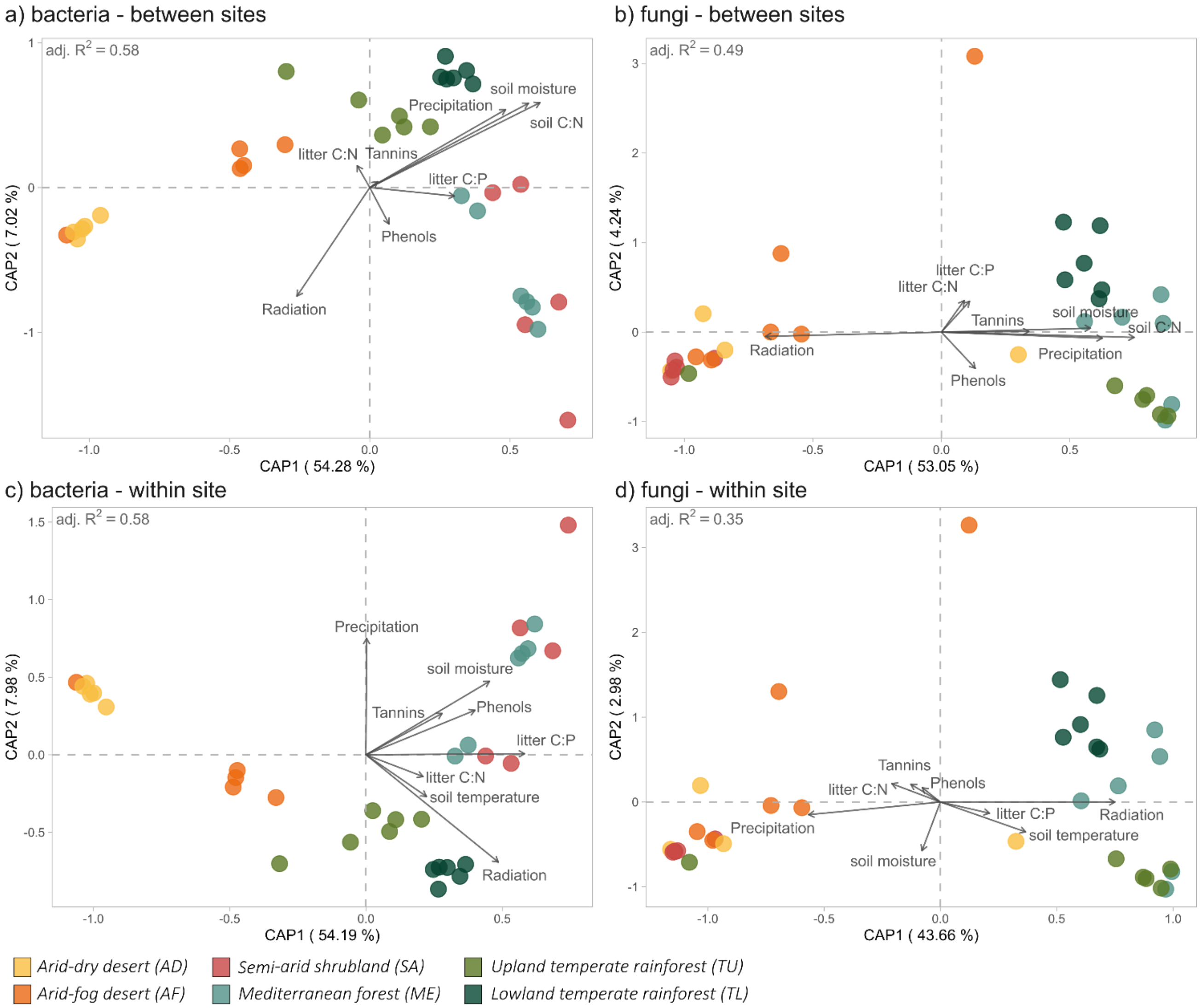


Figure S2: Distance-based redundancy analyses of the functional composition of bacterial (left) and fungal (right) litter communities, with environmental explanatory data sets separated into between-site (top row) and within-site (CV, bottom row) differences. All models were significant with 999 permutations.
